# Engineered Skin Microbiome Reduces Mosquito Attraction to Mice

**DOI:** 10.1101/2023.12.20.572663

**Authors:** Feng Liu, Iliano V. Coutinho-Abreu, Robyn Raban, Tam Thuy Dan Nguyen, Alejandra R. Dimas, Joseph A. Merriman, Omar S. Akbari

## Abstract

The skin microbiome plays a pivotal role in the production of attractive cues detected by mosquitoes. Here we leveraged recent advances in genetic engineering to significantly reduce the production of L-(+)-lactic acid as a strategy to reduce mosquito attraction to the highly prominent skin commensals *Staphylococcus epidermidis* and *Corynebacterium amycolatum*. Engraftment of these engineered bacteria onto the skin of mice reduced mosquito attraction and feeding for up to 11 uninterrupted days, which is considerably longer than the several hours of protection conferred by the leading chemical repellent DEET. Taken together, our findings demonstrate engineering the skin microbiome to reduce attractive volatiles represents an innovative untapped strategy to reduce vector attraction, preventing bites, and pathogen transmission setting the stage for new classes of long-lasting microbiome-based repellent products.

**One-Sentence Summary:** Modified microbes make skin less attractive to mosquitoes

## Main Text

Mosquitoes are responsible for the transmission of a variety of deadly human pathogens, including malaria, West Nile, dengue, yellow fever, and Zika viruses. In 2016, more than 200 million cases of human malaria occurred worldwide (*1*). While it is still considered to be underestimated, the annual dengue cases have increased from approximately half a million in 2000 to 4.2 million in 2019, according to reports from the World Health Organization (*2*).

Current topical repellents are potent inhibitors of mosquito attraction (>90% reduction using DEET) (*3*). However, an important limitation of these repellents is their very short window of protection (*4*), placing individuals at risk of exposure.

Female mosquitoes ingest host blood required for egg development, through which the pathogens are acquired and transmitted by mosquitoes to humans (*1, 5*). Mosquitoes rely on their acute olfactory system to detect volatiles, including CO_2_, L-(+)-lactic acid, and other specific odors to locate their hosts (*6, 7*). Particularly, volatiles emanating from vertebrate skin play essential roles in mosquito host seeking. Indeed, CO_2_ is considered a synergist of skin volatiles, causing stronger behavioral responses in host-seeking mosquitoes than the volatiles alone (*8*–*10*). Volatile compounds inform the mosquito about the quality (*11*) as well as identity (*12*) of the host, being responsible for the host specificity of a range of mosquitoes. While several host-derived compounds affecting mosquito host seeking have been described, L-(+)-lactic acid remains one of the most prominent mosquito attractive volatiles from human emanation, synergizing with CO_2_ in both laboratory and field applications (*8, 13*). It was only recently discovered that many of these compounds, including L-(+)-lactic acid, are of bacterial origin (*14, 15*).

*Staphylococci* and *Corynebacteria* are amongst the most abundant bacterial species found on human skin and primarily found in dry, moist, and sebaceous sites across the body (*16*), providing a large surface area to influence mosquito olfaction. *Anopheles* mosquitoes showed high attraction to the scent of diverse human skin bacteria, including *Corynebacterium sp. (17)*. Additionally, out of the 15 volatiles collected from cultures of human foot microbes, five are also components of the volatile bouquet produced by *S. epidermidis* (*18)*. Both of these bacteria species have been profiled for the production of multiple carboxylic acids specific to the human skin that drive mosquito host seeking behavior (*19*). *Staphylococcus epidermidis (S. epidermidis)* is one of 31 “core” members of the human skin microbiome (*20*), and until recently, has been notoriously difficult to genetically engineer. L-lactate dehydrogenase genes (*l-ldh*) have been identified in both *S. epidermidis* and *C. amycolatum* genomes, indicating a potential to produce L-(+)-lactic acid during growth. We postulated an alternative approach to preventing mosquito bites is the genetic modification of the human skin bacteria to reduce or eliminate the secretion of mosquito attractive odorants (*6, 7*).

To validate the function of *l-ldh* gene of human skin bacteria *S. epidermidis* and *C. amycolatum* in generating L-(+)-lactic acid and attracting mosquitoes, we created *l-ldh* null mutants (Δ*l-ldh)* of both bacteria species. These engineered strains were then tested for attractiveness in a culture-based high throughput olfactometer assay and in the context of mouse colonization for up to 14 days. We confirmed the critical role of *l-ldh* gene of *S. epidermidis* and *C. amycolatum* in their mosquito attraction *in vitro* and *in vivo*. Moreover, we validate the efficacy of these mutant human skin bacteria in reducing mosquito landing and biting using the two-choice non-contact assay and three-choice contact assay. Together, our findings demonstrate the importance of the *l-ldh* gene of human skin bacteria in augmenting the host-seeking process of mosquitoes and support the intriguing perspective of a “living” and long-lasting engineered microbiome-based mosquito repellent.

## Results

### Skin bacteria deficient in L-(+)-lactic acid production are less attractive to mosquitoes

To determine whether engineered skin bacteria could reduce mosquito attraction, we first developed strains of *S. epidermidis* and *C. amycolatum* that were deficient in L-(+)-lactic acid production. To do this, we generated L-lactate dehydrogenase gene deletions in *S. epidermidis* (Suppl. Fig S1A and B) and *C. amycolatum* (Suppl. Fig S1C and D), giving rise to *S. epidermidis* Δ*l-ldh* and *C. amycolatum* Δ*l-ldh*. Neither strain exhibited significant growth defects compared to the parental strain (Suppl. Fig S2A and B). Importantly, both Δ*l-ldh* strains also demonstrated a significant reduction in L-lactate production (Suppl. Fig S2C and D).

As L-(+)-lactic acid, along with carbon dioxide, triggers mosquito short range attraction (*8, 13*) and landing (*19*) behaviors, we implemented a 4-port olfactometer (*21*) to assay the attractive/repellent potential of *S. epidermidis* and *C. amycolatum*, and their respective Δ*l-ldh* counterparts (Fig. 1A). In these experiments, *S. epidermidis* Δ*l-ldh* exhibited repellency to three genera of mosquito species including *Aedes aegypti* (54.2% repellency, Fig 1B), *Culex quinquefasciatus* (21.7% repellency, Fig 1C), and *Anopheles gambiae* (55.9% repellency, Fig 1D) as compared to wildtype (WT). To further validate these findings, we assessed the attraction of the mosquito *Ae. aegypti* to *C. amycolatum* Δ*l-ldh* and showed that these mosquitoes were less attracted to the scent of Δ*l-ldh* as compared to WT cultures (77.1% repellency, Fig 1D). These results demonstrate that, as expected, cultures of the human skin commensals, *S. epidermidis* and *C. amycolatum*, are attractive to *Ae. aegypti, Cx. quinquefasciatus*, and *An. gambiae*. Furthermore, this attraction is significantly reduced when exposed to bacteria engineered to eliminate the production of L-(+)-lactic acid (Δ*l-ldh*).

**Fig. 1.**
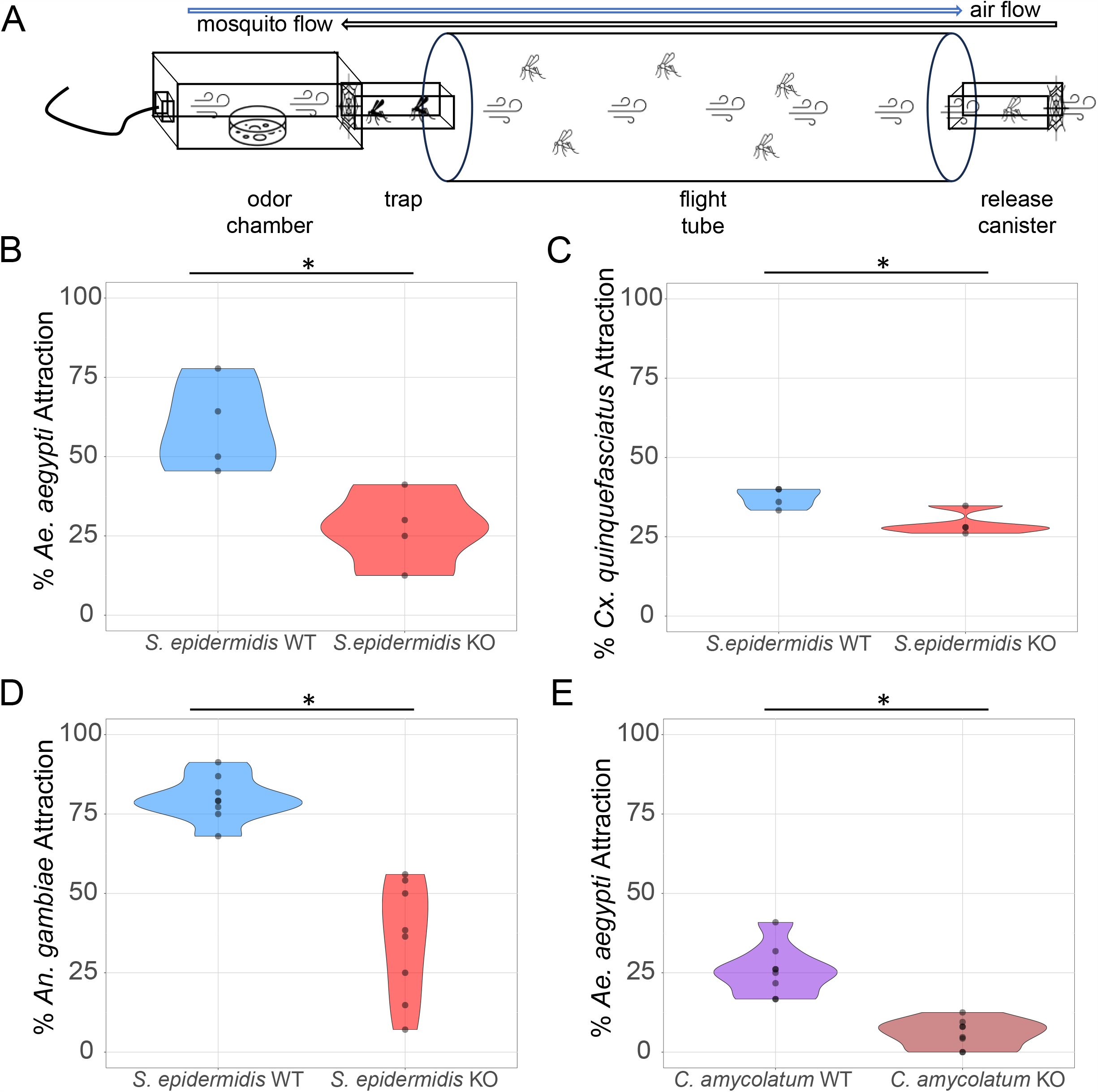
Mosquito attraction to the scent of human skin bacteria cultures in a 4-port olfactometer. (**A)** Schematic of one lane of the 4-port olfactometer. (**B-D**) Attraction to the scent of *Staphylococcus epidermidis* wild type (WT) and L-(+)-lactic acid knockout Δ*l-ldh* (KO) cultures by the mosquitoes *Aedes aegypti* (B), *Culex quinquefasciatus* (C), and *Anopheles gambiae* (D). **(E)** Attraction of the *Ae. aegypti* to the scent of cultures of *Corynebacterium amycolatum* wild type (WT) and Δ*l-ldh* (KO). n = 4-8 biological replicates as represented by each dot. (*) p < 0.05.

### *S. epidermidis* deficient in L-(+)-lactic acid production makes mice less attractive to mosquitoes for multiple days

Mosquito behavior in the context of all host cues (heat, CO_2_, breath, skin odors) is more complex than the strictly *in vitro* olfactometer testing behavior. Therefore to capture these cues, we leveraged a 2-choice non-contact behavioral assay (*19*) to compare mosquito attraction to mice colonized with *S. epidermidis* or *S. epidermidis* Δ*l-ldh* or mice coated with growth media (BHI). *S. epidermidis* cultures were applied onto the skin of mice for three consecutive days (Fig. 2A), and *Ae. aegypti* attraction to these mice was compared to mice treated with culture (BHI) media alone (Fig. 2B-C). Mice engrafted with *S. epidermidis* exhibited greater attraction to mosquitoes compared to BHI-treated mice after 1 (30.5% attraction, Fig. 2D), 3 (84.3% attraction, Fig. 2E), 7 (86.9% attraction, Fig. 2F), and 14 (84.5% attraction, Fig. 2G) days of skin treatment. Further supporting the role of *S. epidermidis* in this increased attraction, *S. epidermidis* was detected at 1 and 14 days after treatment via PCR (Suppl. Fig. 3A and B).

**Fig. 2.**
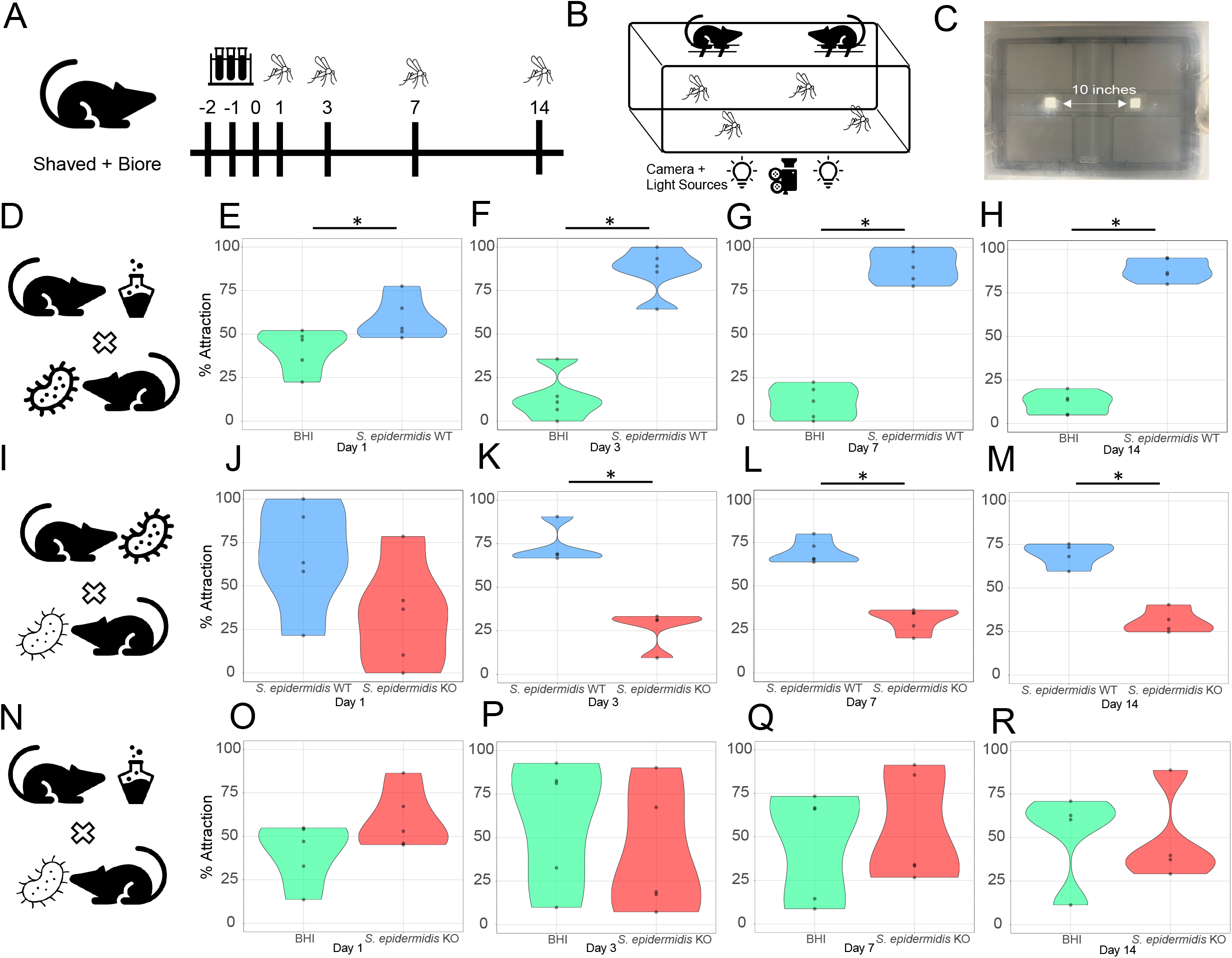
Attraction of the mosquito *Aedes aegypti* to the scent of mice treated with *Staphylococcus epidermidis* wild type strain (WT), L-(+)-lactic acid knockout Δ*l-ldh* strain (KO), and culture media (BHI), in a 2-choice non-contact assay. (**A)** Schematic diagram of mouse skin treatment for skin bacteria engraftment and BHI media coating. (**B)** Diagram of the 2-choice behavioral arena. **C**. Picture depicting the lid of the behavioral arena, highlighting the two windows from which the mosquitoes sense the mice scent. (**D-H)**. Mosquito attraction to mice coated with (**D)** BHI media or engrafted with *S. epidermidis* WT on days 1 (**E**), 3 (**F**), 7 (**G**), and 14 (**H**) after mouse skin treatment. (**I-M)** Mosquito attraction to mice treated with either (**I)** WT or Δ*l-ldh* (KO) strains on days 1 (**J**), 3 (**K**), 7 (**L**), and 14 (**M**) after mouse skin treatment. **N-R**. Mosquito attraction to mice treated with either (**N)** BHI media or Δ*l-ldh* (KO) strain on days 1 (**O**), 3 (**P**), 7 (**Q**), and 14 (**R**) after mouse skin treatment. n = 4-5 biological replicates as represented by each dot. (*) p < 0.05.

**Fig. 3.**
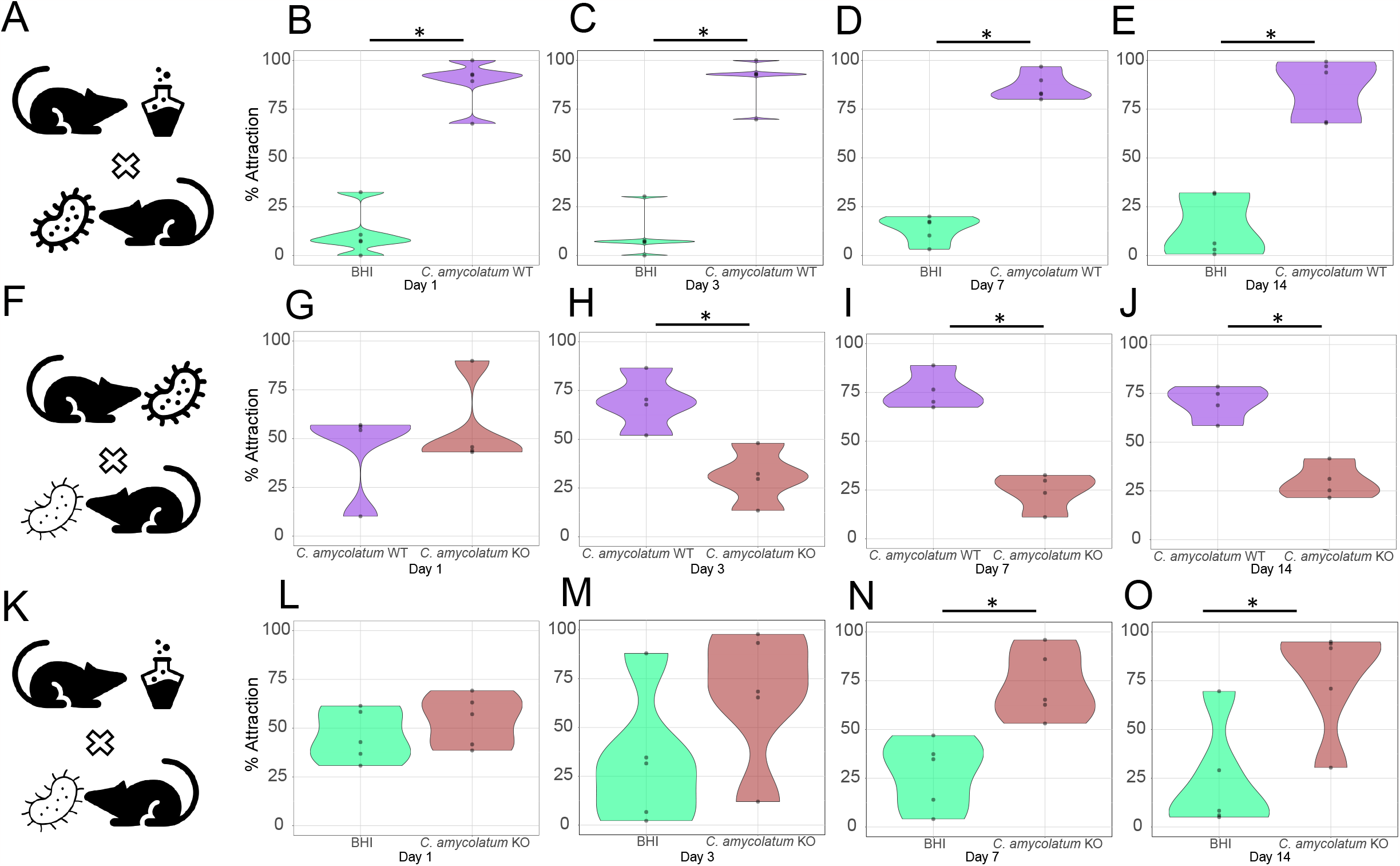
*Aedes aegypti* attraction to *Corynebacterium amycolatum* wild type strain (WT), L-(+)-lactic acid knockout strain Δ*l-ldh* strain (KO), and culture media (BHI) treated mice in a 2-choice non-contact assay. (**A-E)** Mosquito attraction to mice treated with BHI media or wild type (WT) cultures (**A**) after 1 (**B**), 3 (**C**), 7 (**D**), and 14 (**E**) days. (**F-I)** Mosquito attraction to mice treated with WT or Δ*l-ldh* (KO) cultures (**F**) after 1 (**G**), 3 (**H**), 7 (**I**), and 14 (**J**) days. (**K-O)** Mosquito attraction to mice treated with BHI media or Δ*l-ldh* (KO) cultures (**K**) after 1 (**L**), 3 (**M**), 7 (**N**), and 14 (**O**) days. n = 4-5 biological replicates as represented by each dot. (*) p < 0.05.

We next assessed if mice colonized with *S. epidermidis* Δ*l-ldh* led to reduced mosquito attraction compared to the WT strain. Apart from day 1 (Fig 2H), *S. epidermidis* Δ*l-ldh* treated mice showed repellency on days 3 (64.4% repellency, Fig 2I), 7 (56.2% repellency, Fig. 2J), and 14 (55.3% repellency, Fig 2K) after colonization as compared to mice colonized with the WT strain. *S. epidermidis* Δ*l-ldh* may produce D-(+)-lactic acid (*d-ldh* **SERP2087**), however, as it contains a putative lactate racemase gene (ATM22_10565) that may convertD-lactate to L-lactate. To address the possible impact of this alternative L-lactate production pathway on mosquito attraction, we tested mosquito attraction to mice colonized with *S. epidermidis* Δ*l-ldh* and compared to mice treated with BHI media (Figs 2L-O). Mosquitoes did not exhibit differential attraction towards *l-ldh* knockout treated-mice at any time point (Figs 2L-O), despite similar levels of colonization between *S. epidermidis* Δ*l-ldh* and WT colonized mice (Suppl. Fig 3B). Together, these findings demonstrate that *S. epidermidis* Δ*l-ldh* abrogated mosquito attraction to mice for 11 uninterrupted days without any residual attractive effect.

### Mosquitoes are less attracted to mice treated with *Corynebacterium amycolatum* deficient in the synthesis of L-(+)-lactic acid

*Corynebacterium spp*. also represents a significant proportion of the human skin microbiome and plays a role in mosquito attraction (*17*). Significant mosquito behavioral changes to mice colonized with *S. epidermidis* Δ*l-ldh* led us to consider if this phenomenon is conserved in another prominent skin commensal, *C. amycolatum*. To this end, following experimental procedures used to test *S. epidermidis* colonized mice, we tested *C. amycolatum* WT or Δ*l-ldh* colonized mice and assessed mosquito attraction in the 2-choice behavioral model (Fig 3).

As observed for *S. epidermidis, C. amycolatum* WT colonized mice evoked much stronger mosquito attraction than BHI media-associated mice on days 1 (91.8% attraction, Fig. 3A), 3 (92.3% attraction, Fig 3B), 7 (79.7% attraction, Fig. 3C), and 14 (82.7% attraction, Fig. 3D) post-association. Next, we compared mosquito preferential attraction to mice treated with *C. amycolatum* Δ*l-ldh* and mice treated with the WT strain (Figs. 3E-H). Analogous to *S. epidermidis* Δ*l-ldh* colonized mice, mice colonized with *C. amycolatum* Δ*l-ldh* showed repellency to female *Ae. aegypti* mosquitoes when compared to WT colonized mice on days 3 (55.4% repellency, Fig. 3F), 7 (68.0% repellency, Fig 3G), and 14 (57.4% repellency, Fig 3H) after skin engraftment. However, *C. amycolatum* still exhibited some attractive residual effects (Figs 3I-L). In trials between mice treated with either BHI media or *C. amycolatum* Δ*l-ldh*, the latter group exhibited significantly greater attraction to *Ae. aegypti* on days 7 (62.2% attraction, Fig. 3K) and 14 (69.1% attraction Fig 3L) after skin colonization. Despite exhibiting residual attractive effects, mice treated with *C. amycolatum* Δ*l-ldh* repelled mosquitoes compared to WT colonized mice.

### Colonization with *S. epidermidis* Δ*l-ldh* reduces mosquito feeding propensity

To elucidate the effect of skin bacteria colonization on mosquito feeding behaviors, we modified the behavioral arena to a 3-choice contact assay (Fig. 4A and B). In this assay, *Ae. aegypti* were exposed to mice treated with *S. epidermidis* WT, *S. epidermidis* Δ*l-ldh*, or BHI media and allowed to choose which one to feed upon. In this 3-choice setup, mosquitoes still displayed repellency (Figs. 4C-G) to mice treated with *S. epidermidis* Δ*l-ldh* compared to the WT strain on days 7 (64.2% repellency, Fig. 4F) and 14 (64.6% repellency, Fig. 4G) after skin engraftment. Even though mice treated with *S. epidermidis* Δ*l-ldh* presented greater attraction than BHI media treated ones on day 3 (46.2% attraction, Fig. 4E), that residual attractive effect was lost on day 7 (Fig. 4F) and turned into repellency on day 14 (42.1% repellency, Fig. 4G).

**Fig. 4.**
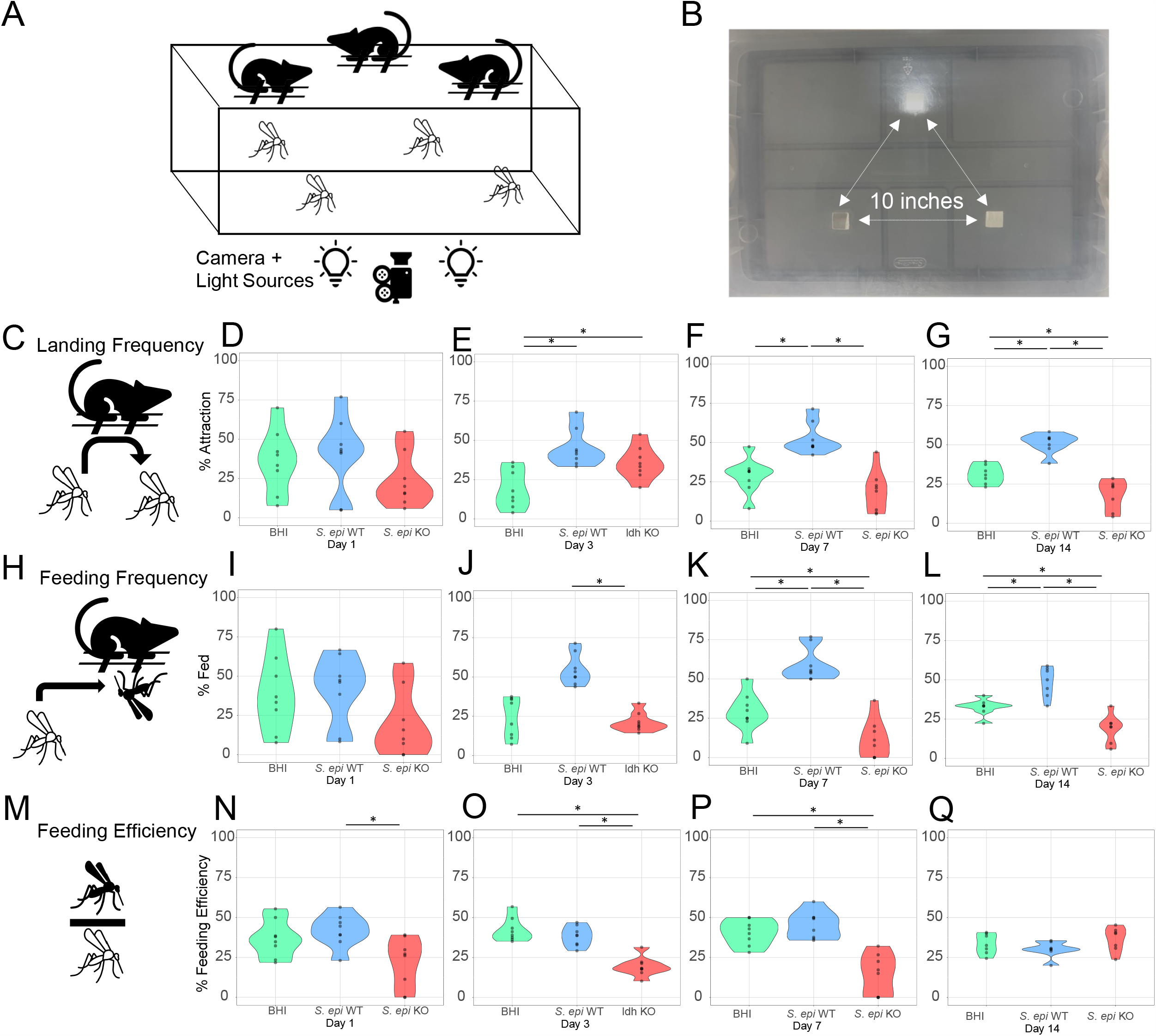
Attraction and feeding behavior of the mosquito *Aedes aegypti* exposed to mice treated with *Staphylococcus epidermidis* wild type strain (WT), L-(+)-lactic acid knockout Δ*l-ldh* strain (KO), and culture media (BHI) in a 3-choice contact assay. (**A)** Diagram of the 3-choice behavioral arena. (**B)** Image depicting the lid of the behavioral arena, highlighting the three equidistant windows from which the mosquitoes sense the scent and feed upon the mice. (**C-G)** Mosquito attraction (landing frequency, **C**) to mice on days 1 (**D**), 3 (**E**), 7 (**F**), and 14 (**G**) after mouse skin treatment. (**H-L)** Mosquito feeding (feeding frequency, **H**) on mice after 1 (**I**), 3 (**J**), 7 (**K**), and 14 (**L**) days of skin treatment. (**M-Q)** Mosquito feeding efficiency (% # fed/# attracted, **M**) on days 1 (**N**), 3 (**O**), 7 (**P**), and 14 (**Q**) after mouse skin treatment. n = 7-8 biological replicates as represented by each dot. (*) p(adjusted) < 0.05.

Using this model, we assessed whether this *S. epidermidis* Δ*l-ldh* also affects *Ae. aegypti* feeding abilities (Figs. 4H-L). By determining the number of mosquitoes that engorged from each mouse treated with *S. epidermidis* WT, *S. epidermidis* Δ*l-ldh*, or BHI medium, we observed that fewer mosquitoes took blood meals from mice treated with the Δ*l-ldh* strain compared to WT treated mice on days 3 (61.2% deterrence, Fig. 4J), 7 (80.6% deterrence, Fig 4K), and 14 (60.7% deterrence, Fig. 4L). Furthermore, similar numbers of mosquitoes on days 1 and 3 (Figs. 4I-J) and fewer mosquitoes on days 7 and 14 (41.5-60.7% deterrence range, Figs. 4K-L) after colonization took blood meals from mice treated with the Δ*l-ldh* strain compared to BHI media-treated mice.

Besides reducing mosquito landing and feeding for multiple days, it is important to assess if mice treated with *S. epidermidis* Δ*l-ldh* strain affect mosquito feeding efficiency (Figs. 4M-Q): the ratio of the number of mosquitoes that actually fed upon each mouse out of the mosquitoes that landed on them (Fig. 4M). Feeding efficiencies were reduced upon mice colonized with *S. epidermidis* Δ*l-ldh* when compared to the WT treated on days 1 (48.5% reduced feeding efficiency, Fig. 4N), 3 (49.7% reduced feeding efficiency, Fig 4O), and 7 (64.7% reduced feeding efficiency, Fig 4P) after skin treatment. Mice treated with *S. epidermidis* Δ*l-ldh* displayed reduced feeding efficiency to mosquitoes on days 3 (54.4% reduced feeding efficiency, Fig. 4O) and 7 (61.1% reduced feeding efficiency, Fig. 4P) compared to mice associated with BHI media. Altogether, these findings indicate that *S. epidermidis* Δ*l-ldh* evokes a feeding deterrent behavior in mosquitoes that lasts at least one week.

## Discussion

Mosquito host seeking behavior is mediated by the volatiles released by the human breath and by the resident skin bacteria (*6*). Among those, carbon dioxide mediates mosquito activation and hierarchically interacts with L-(+)-lactic acid, ammonia, and other short and middle chain carboxylic acids to induce strong host attraction (*9, 10, 13, 22*). The absence of components like L-(+)-lactic acid and ammonia in an odor blend significantly reduces the ability of mosquitoes to perform odor-mediated attraction and landing (*8*–*10, 19*).

DEET remains the gold standard in topically applied repellents (> 90% reduction in attraction)(*3*). However, the need for constant re-application (within hours (*4*)) leads to logistical issues and is cost prohibitive in malaria endemic regions worldwide. Amongst novel strategies to prevent mosquito bites, the reengineering of the human skin microbiome to produce repellents and/or reduced levels of attractive odorants may be realized as a more stable and long-lasting solution (*6, 23*). As L-(+)-lactic acid and ammonia are key odorants to gate mosquito human seeking behavior, reducing the production of such odorants by the skin microbiome might result in an effective strategy to reduce mosquito bites and pathogen transmission.

In this study, we engineered two common human skin commensals, *S. epidermidis* and *C. amycolatum*, and significantly decreased their production of L-(+)-lactic acid through deletion of the L-lactate dehydrogenase (Δ*l-ldh*) gene. We also demonstrated that mosquitoes are less attracted to the scent of the Δ*l-ldh* strains than to WT *in vitro*. These findings prove that skin-bacterium deprived of the ability to produce L-(+)-lactic acid is key to reducing mosquito attraction, despite these bacteria producing other human skin derived odorants (*18*).

Building upon these findings, we tested the effects of these strains on mosquito behavior after colonizing a mouse model in 2-choice non-contact or 3-choice contact assays. Upon engraftment, WT strains increased mosquito attraction for two weeks. In contrast, mice colonized with Δ*l-ldh* counterparts are less attractive to mosquitoes for 11 consecutive days with little to no residual attractive effect. Mosquito feeding desire is also reduced for seven consecutive days. While not equivalent to protection conferred by DEET or picaridin (90-100%) (*3*), we observed significant repellency (55.3-68%) and deterrence (60.7%-80.6%) mediated by Δ*l-ldh* skin bacteria. Notably, this protection lasted 7-11 days post application, whereas DEET/picaridin is only effective for a few hours, requiring constant reapplication (*3, 4*).

This study demonstrates the potent effect of human skin commensal derived L-(+)-lactic acid on mosquito attraction and feeding efficiency for 7-11 days. As an approach, living mosquito repellents benefit from 1) durable, self-replicating protection (no lapse in protection concerns), 2) low logistical burden, and 3) significantly cheaper lifetime protection. Provided genetically engineered (Δ*l-ldh)* skin bacteria can grant effective and long lasting repellency to the human skin, this novel strategy could be used alone or in combination with topical application of synthetic repellents to reduce vectorial capacity and provide long-lasting skin protection from mosquito bites, pathogen transmission, and mosquito-borne diseases.

## Materials and Methods

### Mosquito rearing

*Aedes aegypti* Liverpool, *C. quinquefasciatus* wild type S-strain, and *A. gambiae* G3 strain mosquitoes were raised at 28.0°C and 70% relative humidity (12 hr light/dark cycle). Larvae were fed with ground fish food (TetraMin Tropical Flakes, Tetra Werke, Melle, Germany), and the *A. gambiae* diet was supplemented with 2% beef liver powder (NOW, Bloomingdale, IL, USA). Adults were provided 0.3 M aqueous sucrose *ad libitum*. Adult females were blood fed three to five days after emergence upon anesthetized mice. All animals were handled in accordance with the Guide for the Care and Use of Laboratory Animals as recommended by the National Institutes of Health and supervised by the local Institutional Animal Care and Use Committee (S17187) and the Animal Care and Use Review Office (ACURO) protocol DARPA-9729.

### Genetic engineering of skin bacteria

*Staphylococcus epidermidis* NIHLM087 (ATM22_01530) was obtained from the NIH (*24*). *Corynebacterium amycolatum* ATCC 49368 (16165_RS07200) was purchased from ATCC: The Global Resource Center. All bacterial strains were grown in Difco Brain Heart Infusion (BHI) media (BD 237200) at 37°C with shaking in which *Corynebacteria* species were cultured in BHI media supplemented with 1% Tween-80 (BHIT). Prior to liquid growth, individual colonies were cultured overnight on respective BHI (for *Staphylococcus* strains) or BHIT (for *Corynebacterium* strains) agar plates. To delete *l-ldh* genes in *S. epidermidis* and *C. amycolatum*, ∼1,000 bp directly upstream and downstream of the *l-ldh* gene were amplified for each strain and the PCR products were cloned into the pIMAY (for *S. epidermidis* NIHLM087; Addgene Plasmid #68939) and pJSC232 (for *C. amycolatum* ATCC 49368) temperature sensitive vectors using Gibson assembly. Transformation of the species-specific deletion vector was performed as described in (*25*). To generate electrocompetent cells, overnight *S. epidermidis* grown in BHI supplemented with 0.5 M sorbitol (Sigma) (BHIS) and *C. amycolatum* grown in BHIT supplemented with 0.5 M sorbitol (BHIST) were back-diluted to an optical density of 0.15 for *S. epidermidis* and 0.3 for *C. amycolatum*. Back-diluted cultures were placed on ice once they reached an OD of 0.7 for *S. epidermidis* and 1.2 for *C. amycolatum* and pelleted at 3,500g for 10 min at 4°C. Spun-down cultures were resuspended in equal volume with 10% ice-cold glycerol followed by five 10% glycerol washes. After the last wash, cells were suspended in 100 μL of 10% ice-cold glycerol to use for electroporation.

Approximately 1 μg DNA of plasmid isolated from DC10B and TOP10 *Escherichia coli* was added respectively to 100 μL of competent *S. epidermidis* and *C. amycolatum*. For methods specific to *S. epidermidis*, cells and plasmids in 10% glycerol were first heat-shocked at 56°C for 2 min followed by immediate electroporation in 0.1-cm cuvette (Bio-Rad) using electroporation program of 2.5 kV and time constant of 2.3-2.5 ms on the Bio-Rad Micropulser. Electroporated cells were then transferred to 3 mL of pre-warmed BHIS and recovered at 37°C for 3 hours prior to plating on BHIS plates with appropriate antibiotics. For methods specific to *C. amycolatum*, plasmid was electroporated into competent cells in 0.2-cm cuvette with 2.5 kV and time constant of 4 ms and immediately heat-shocked in BHIST that was previously pre-warmed at 46°C for 6 min then recovered at 37°C for 3 hours prior to plating on BHIST plates with appropriate antibiotics.

To create targeted deletion of *l-ldh* in *S. epidermidis* LM087 and *C. amycolatum* ATCC 49368 (*25*), once species-specific deletion plasmid was electroporated into each strain, transformed cells were selected on 10 μg/mL chloramphenicol (for *S. epidermidis*) and 25 μg/mL kanamycin (for *C. amycolatum*) for single chromosomal crossover, followed by selection solid medium containing 1 μg/mL anhydrotetracycline (for *S. epidermidis*) and 10 μg/mL sucrose (for *C. amycolatum*), which produces either complete gene deletions or wild type bacteria revertants. Gene deletions are verified by PCR and Sanger sequencing.

All the primers used in the above experiments are listed in (Suppl. Table 1). *S. epidermidis* LM087 Δ*l-ldh* was fully sequenced on an Illumina NextSeq500 and compared to Wt LM087 using Snippy (https://github.com/tseemann/snippy) and found to have 100% coverage with no SNPs or indels, except for the absence of 855bp found within the L-ldh gene.

### 4-port olfactometer assay

Mosquito behavioral assays were performed with a modified 4-port high-throughput olfactometer (*21*) at 27°C and 80% relative humidity. The screens of the olfactometer’s traps were removed to allow mosquitoes to make a choice between staying close or moving far away from the odor source. Twenty female mosquitoes were transferred to the releasing canisters and starved for 5-8 hours without water. Purified air was pumped into the system at 24,367 mL/min rate, whereas pure CO_2_ was flown at 254 mL/min (final concentration per lane ∼ 1,500-2000 ppm). Mosquitoes were exposed to air only for 10 minutes, when bacterial cultures (1,000μl) were placed in the odor chamber onto 47mm plastic petri dishes (Fisher Scientific, Hampton, NH), and CO_2_ gauge was switched on. The gates of the releasing canisters were open, and the behavioral assays were carried out for 20 min. Then, both the releasing canister and the trap gates were closed, and the number of mosquitoes in the releasing canisters, flight tubes, and traps were scored.

### Mouse engraftment procedure

Wild type and Δ*l-ldh S. epidermidis* and *C. amycolatum* strains were inoculated onto BHI plates from frozen stocks. Single colonies were picked to individually inoculate 4 mL BHI media in 15 mL culture vials for overnight growth before engraftment. OD_600_ values of WT and Δ*l-ldh* cultures were measured with Nanodrop and normalized to 2.0 with BHI media. For skin engraftment, 6-to-8-week-old female C57BL/6 mice were purchased from Jackson Laboratory (Jax). Twenty anesthetized mice were shaved in either the abdomen or flank region. Native microbes were removed using Biore Deep Cleansing Pore Strips (Biore, Cincinnati, OH) following the manufacturer’s instruction (10 minutes application). Bacteria strains were engrafted on the shaved abdomen of mice by dipping a swab (ESwab Collection Kit, Becton, Dickinson, and Company, Franklin Lakes, NJ) into the bacterium culture and swabbing the exposed skin of the mouse abdomen 15 times for three consecutive days. The same procedure was conducted with bacterial BHI medium, which served as a control for the experiment.

### 2-choice non-contact mouse assay

Plastic containers (50cm x 30cm x 15cm, Hefty) were modified by cutting two square windows (1 x 1 inch) on the lid 30 cm (10 inches) apart. Square polyester meshes (2 x 2 inches) were used to cover the windows from the inner side to prevent mosquito escape. Custom-made plastic frames (1/16” thick, gates used for the 4-port olfactometer, (*21*) with small holes were placed on the outer side of the windows to create a short distance between mouse skin and mosquito, preventing the mosquitoes from having physical contact with the mice. Twenty mosquitoes were introduced into the box using a mouth aspirator (John Hock, Gainesville, FL) before the trials. Two mice (BHI-treated versus WT bacteria-treated, WT bacteria versus Δ*l-ldh* bacteria-treated, or BHI-treated versus *ldh* KO bacterium-treated) were placed on the top of plastic frames with their shaved abdomen facing the experimental cage. Host-seeking activity of mosquitoes was recorded for ten minutes. Mice with different treatments were switched between windows across trials to prevent positional bias. The videos were processed by manually counting the landing frequency of mosquitoes.

### 3-choice contact mouse assay

The lids of similar arenas used in the 2-choice assays were modified by cutting three square windows (1 x 1 inch) 10 inches apart in a triangular shape. Unlike the two-choice non-contact assay, no mesh was used to cover the window, and no plastic frame was placed on top of the window, enabling the mosquito to contact the mouse skin and initiate blood feeding. Twenty female mosquitoes were introduced into the arena with a mouth aspirator. Three mice treated with BHI media, WT *S. epidermidis* or Δ*l-ldh S. epidermidis* were placed on the windows with their shaved abdomen facing the arena. The host-seeking and blood feeding activities of mosquitoes were recorded for ten minutes. To eliminate any potential position effect, we interchanged the position of the mice with different treatments across replicates. The videos are further processed by manually counting the mosquito landing and feeding.

### Video recording of behavioral activity

For the 2- and 3-choice assays, videos of mosquito activity were recorded with an iPhone X at 30 fps. Videos were analyzed by blinded, visual counting.

### Behavior apparatus cleaning

The 4-port olfactometer parts were soaked overnight (small parts) or washed thoroughly (flight tubes) with scent-free laundry detergent (Seventh Generation, free & clear) and rinsed with tap water thoroughly. For the 2- and 3-choice arenas, all the parts in contact with the mice were washed with the same detergent.

### Statistical analyses

Graphs and statistical analyses were performed with the R software, using the ggplot2 package. For all experiments, the number of mosquitoes landing and feeding on each mouse and the number of mosquitoes caught by the traps were transformed into percentages to normalize mosquito participation variability across experimental replicates (Supplemental Table S1).

Percent attraction was calculated as (1 - number of landing events on control mouse/number of landing events on treated mouse) x 100. Percent repellency was calculated as (1 - number landing events on treated mouse/number of landing events of control mouse) x 100. Percent deterrence was calculated as (1 - number feeding events on treated mouse/number of landing events of control mouse) x 100. Shapiro-Wilk normality test was used to assess whether the data fit a normal distribution. For pairwise comparisons, either the Welsh t-test or Wilcoxon rank sum test were used. For multiple comparisons, either ANOVA or Kruskal-Wallis’s rank sum test were applied. These tests were followed by post-hoc analyses using Tukey multiple comparisons of means and Wilcoxon rank sum test, respectively. p-values were adjusted (p-adjusted) for multiple comparisons using the Benjamini-Hochberg procedure. The raw data and R codes are provided in the (Supplemental Table S1).

## Acknowledgments

We thank Judy Ishikawa and Ava Stevenson for helping with mosquito husbandry. We thank Michael Fischbach for helping produce the engineered bacteria used in this study. We thank Allison Weakley, Ashley Cabrera, and Xiandong Meng for sequencing and analyzing the *S. epidermidis* Δ*l-ldh* strain for this paper. The views, opinions, and/or findings expressed are those of the authors and should not be interpreted as representing the official views or policies of the U.S. government.

## Author contributions

Conceptualization: OAS

Methodology: FL, ICA, RR, JAM

Investigation: FL, ICA, TN, ARD

Visualization: ICA, FL, TN, ARD, RR

Funding acquisition: OAS

Project administration: RR, JAM

Supervision: OAS, ICA, RR, JM

Writing – original draft: ICA, FL

Writing – review & editing: OAS, RR, JAM

## Data and materials availability

All data are available in the main text or the supplementary materials.

**Fig. S1.**
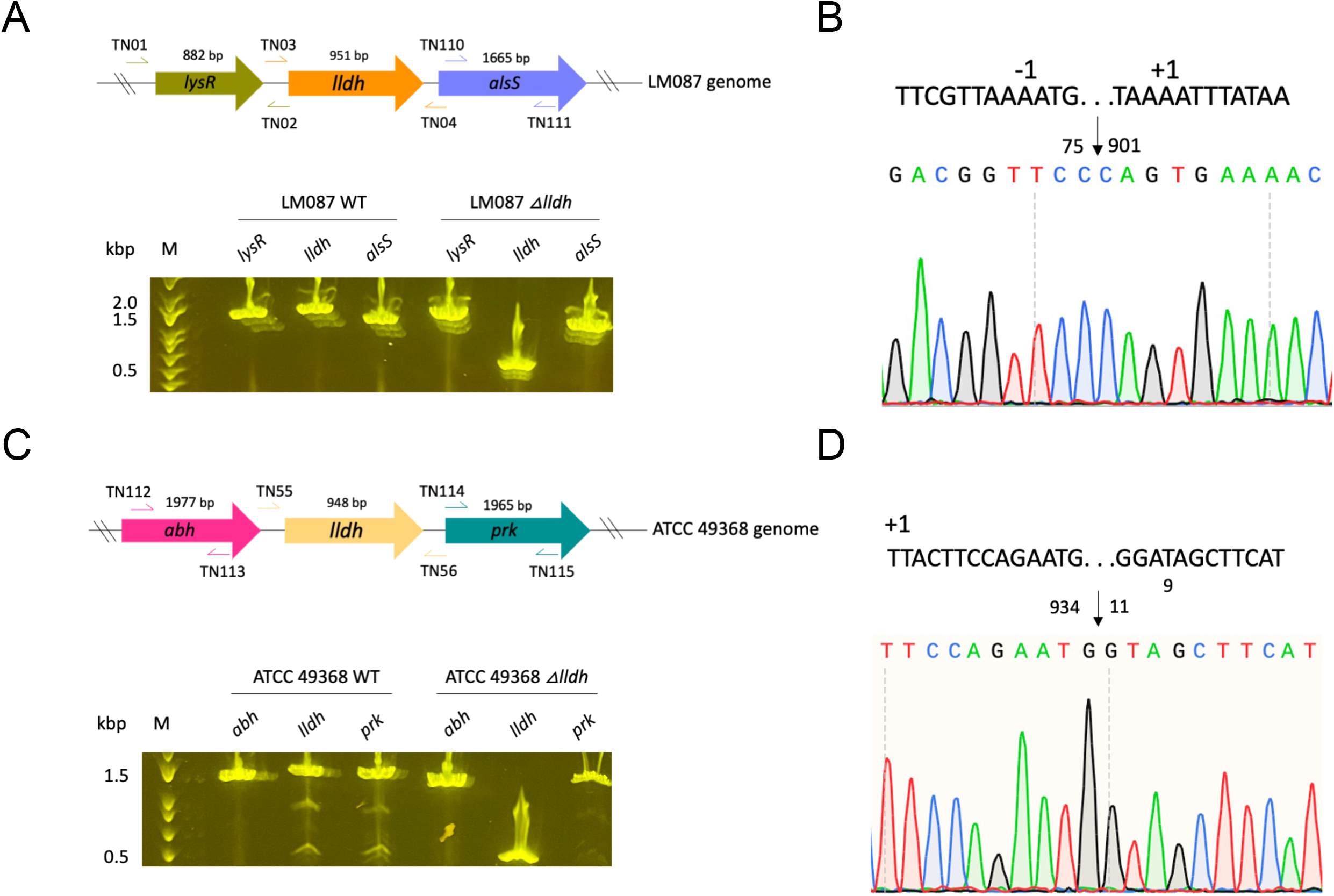
Molecular characterization of the Δ*l-ldh* deletion. (**A**) Schematic drawing of gene loci and primer sites of the *Staphylococcus epidermidis* gene *l-ldh*, and nearby genes *lysR* and alsS used as control, on the top. PCR confirmation of *l-ldh* gene deletion in *S. epidermidis* on the bottom. Neither nearby lysR nor alsS genes were affected by the *l-ldh* gene knockout procedure. (**B**) Demonstration of the Δ*l-ldh* mutation in *S. epidermidis* through Sanger sequencing. (**C**) Primer target sites for the *l-ldh* gene, as well as the nearby abh and prk genes, in *Corynebacterium amycolatum* on the top. On the bottom, PCR gel confirmation of the Δ*l-ldh* mutation in *C. amycolatum*. (**D**) Sanger sequencing confirmation of the deletion in the *C. amycolatum l-ldh* gene.

**Fig. S2.**
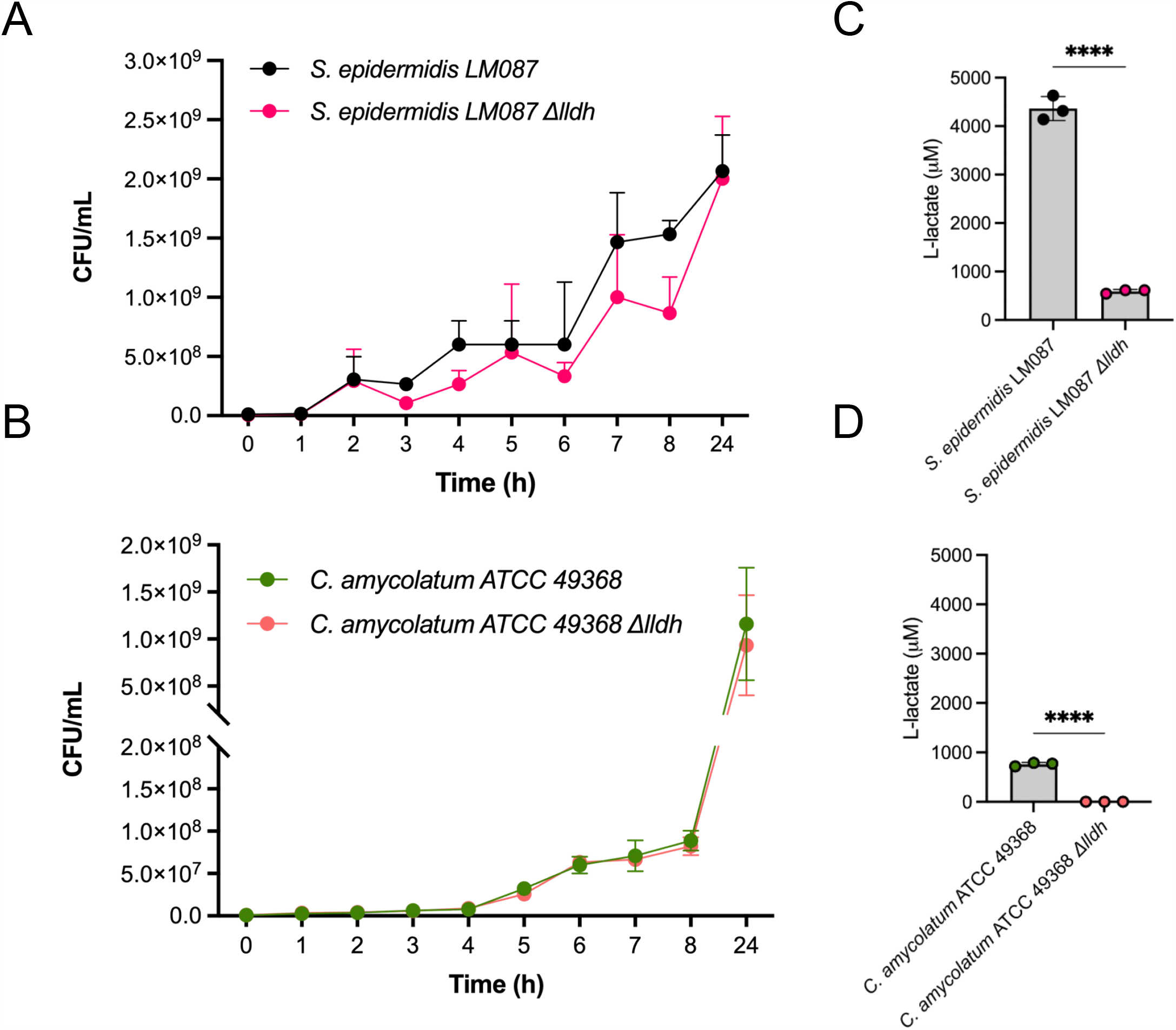
Growth curves and L-(+)-lactic acid production. Growth curves of *Staphylococcus epidermidis* (**A)** and *Corynebacterium amycolatum* **(B)** wild type (WT) and L-(+)-lactic acid knockout Δ*l-ldh* (KO) strains for 24 hours. Colorimetric assay detection of L-(+)-lactic acid in cultures of *S. epidermidis* (**C)** and *C. amycolatum* **(D)** wild type (WT) and L-(+)-lactic acid knockout Δ*l-ldh* (KO) strains. n = 3. (****) p < 0.05.

**Fig. S3.**
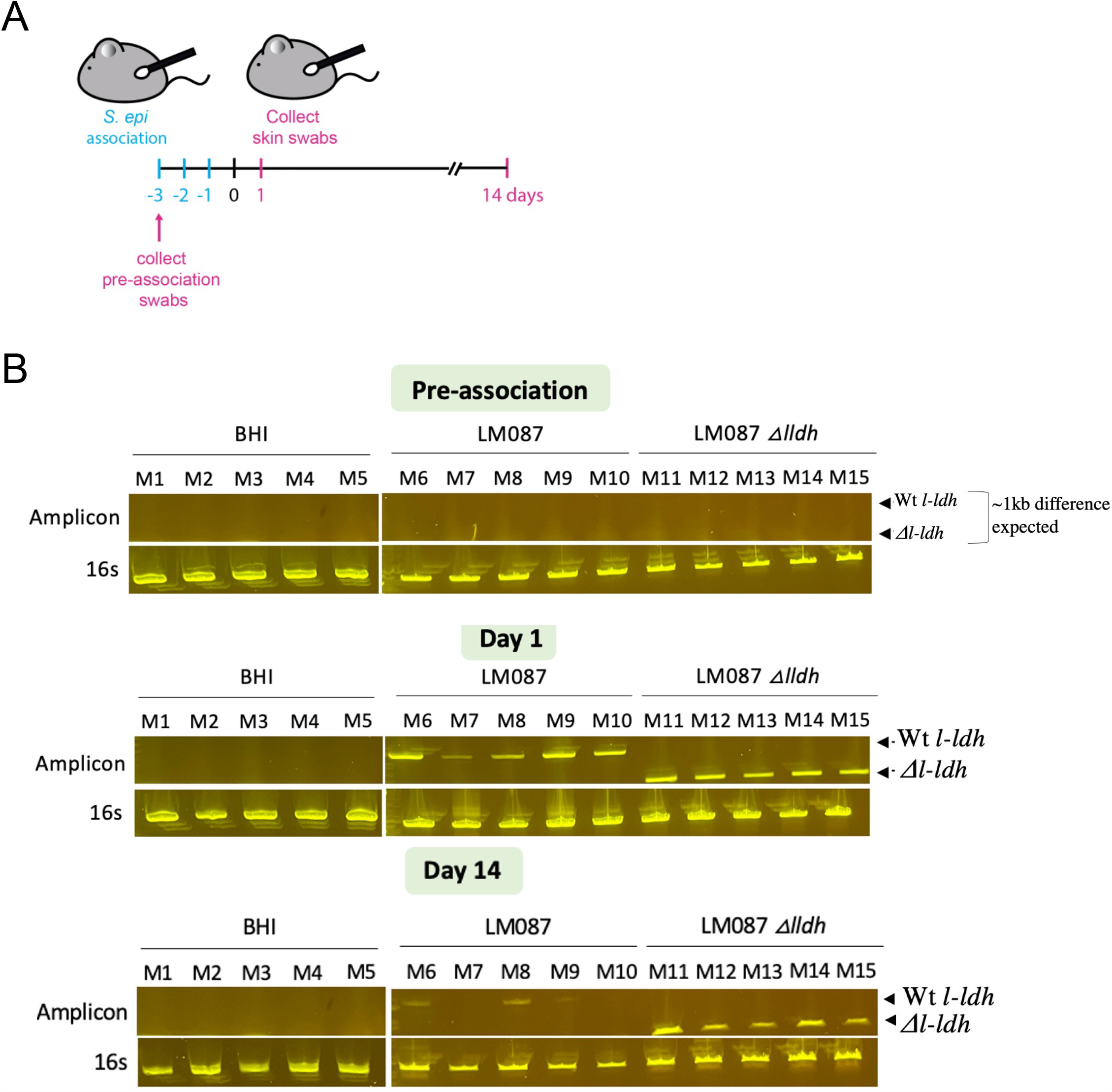
Detection of *Staphylococcus epidermidis* wild type (WT) and L-(+)-lactic acid knockout Δ*l-ldh* (KO) on mice skin swabs. (**A**) Schematic drawing of the timeline used for mouse engraftment with bacteria and swab collection of samples. (**B**) Detection of *S. epidermidis* on skin swabs of mice engrafted with wild type and Δ*l-ldh* strains collected before engraftment (top), 1 day (middle), and 14 days (bottom) after skin treatment.

## References and Notes

1. P. Alonso, A. M. Noor, The global fight against malaria is at crossroads. Lancet. 390, 2532–2534 (2017).

2. World Health Organization, World malaria report 2020: 20 years of global progress and challenges (World Health Organization, 2020).

3. E. Lupi, C. Hatz, P. Schlagenhauf, The efficacy of repellents against Aedes, Anopheles, Culex and Ixodes spp. - a literature review. Travel Med. Infect. Dis. 11, 374–411 (2013).

4. W. S. Leal, The enigmatic reception of DEET - the gold standard of insect repellents. Curr Opin Insect Sci. 6, 93–98 (2014).

5. J. Cox, J. Mota, S. Sukupolvi-Petty, M. S. Diamond, R. Rico-Hesse, Mosquito bite delivery of dengue virus enhances immunogenicity and pathogenesis in humanized mice. J. Virol. 86, 7637–7649 (2012).

6. I. V. Coutinho-Abreu, J. A. Riffell, O. S. Akbari, Human attractive cues and mosquito host-seeking behavior. Trends Parasitol. 38, 246–264 (2022).

7. J. K. Konopka, D. Task, A. Afify, J. Raji, K. Deibel, S. Maguire, R. Lawrence, C. J. Potter, Olfaction in Anopheles mosquitoes. Chem. Senses. 46 (2021), doi:10.1093/chemse/bjab021.

8. F. Acree Jr, R. B. Turner, H. K. Gouck, M. Beroza, N. Smith, L-Lactic acid: a mosquito attractant isolated from humans. Science. 161, 1346–1347 (1968).

9. R. C. Smallegange, Y. T. Qiu, J. J. A. van Loon, W. Takken, Synergism between ammonia, lactic acid and carboxylic acids as kairomones in the host-seeking behaviour of the malaria mosquito Anopheles gambiae sensu stricto (Diptera: Culicidae). Chem. Senses. 30, 145–152 (2005).

10. C. J. McMeniman, R. A. Corfas, B. J. Matthews, S. A. Ritchie, L. B. Vosshall, Multimodal integration of carbon dioxide and other sensory cues drives mosquito attraction to humans. Cell. 156, 1060–1071 (2014).

11. D. Giraldo, S. Rankin-Turner, A. Corver, G. M. Tauxe, A. L. Gao, D. M. Jackson, L. Simubali, C. Book, J. C. Stevenson, P. E. Thuma, R. C. McCoy, A. Gordus, M. M. Mburu, E. Simulundu, C. J. McMeniman, Human scent guides mosquito thermotaxis and host selection under naturalistic conditions. Curr. Biol. 33, 2367–2382.e7 (2023).

12. N. O. Verhulst, A. Umanets, B. T. Weldegergis, J. P. A. Maas, T. M. Visser, M. Dicke, H. Smidt, W. Takken, Do apes smell like humans? The role of skin bacteria and volatiles of primates in mosquito host selection. J. Exp. Biol. 221 (2018), doi:10.1242/jeb.185959.

13. W. R. Mukabana, C. K. Mweresa, B. Otieno, P. Omusula, R. C. Smallegange, J. J. A. van Loon, W. Takken, A novel synthetic odorant blend for trapping of malaria and other African mosquito species. J. Chem. Ecol. 38, 235–244 (2012).

14. N. O. Verhulst, Y. T. Qiu, H. Beijleveld, C. Maliepaard, D. Knights, S. Schulz, D. Berg-Lyons, C. L. Lauber, W. Verduijn, G. W. Haasnoot, R. Mumm, H. J. Bouwmeester, F. H. J. Claas, M. Dicke, J. J. A. van Loon, W. Takken, R. Knight, R. C. Smallegange, Composition of human skin microbiota affects attractiveness to malaria mosquitoes. PLoS One. 6, e28991 (2011).

15. R. C. Smallegange, N. O. Verhulst, W. Takken, Sweaty skin: an invitation to bite? Trends Parasitol. 27, 143–148 (2011).

16. A. L. Byrd, Y. Belkaid, J. A. Segre, The human skin microbiome. Nat. Rev. Microbiol. 16, 143–155 (2018).

17. N. O. Verhulst, R. Andriessen, U. Groenhagen, G. Bukovinszkiné Kiss, S. Schulz, W. Takken, J. J. A. van Loon, G. Schraa, R. C. Smallegange, Differential attraction of malaria mosquitoes to volatile blends produced by human skin bacteria. PLoS One. 5, e15829 (2010).

18. N. O. Verhulst, H. Beijleveld, B. G. Knols, W. Takken, G. Schraa, H. J. Bouwmeester, R. C. Smallegange, Cultured skin microbiota attracts malaria mosquitoes. Malar. J. 8, 302 (2009).

19. I. V. Coutinho-Abreu, O. Jamshidi, R. Raban, K. Atabakhsh, J. A. Merriman, M. A. Fischbach, O. S. Akbari, Identification of human skin microbiome odorants that manipulate mosquito landing behavior. bioRxiv, 2023.08.19.553996 (2023).

20. D. Danko, D. Bezdan, E. E. Afshin, S. Ahsanuddin, C. Bhattacharya, D. J. Butler, K. R. Chng, D. Donnellan, J. Hecht, K. Jackson, K. Kuchin, M. Karasikov, A. Lyons, L. Mak, D. Meleshko, H. Mustafa, B. Mutai, R. Y. Neches, A. Ng, O. Nikolayeva, T. Nikolayeva, E. Png, K. A. Ryon, J. L. Sanchez, H. Shaaban, M. A. Sierra, D. Thomas, B. Young, O. O. Abudayyeh, J. Alicea, M. Bhattacharyya, R. Blekhman, E. Castro-Nallar, A. M. Cañas, A. D. Chatziefthimiou, R. W. Crawford, F. De Filippis, Y. Deng, C. Desnues, E. Dias-Neto, M. Dybwad, E. Elhaik, D. Ercolini, A. Frolova, D. Gankin, J. S. Gootenberg, A. B. Graf, D. C. Green, I. Hajirasouliha, J. J. A. Hastings, M. Hernandez, G. Iraola, S. Jang, A. Kahles, F. J. Kelly, K. Knights, N. C. Kyrpides, P. P. Łabaj, P. K. H. Lee, M. H. Y. Leung, P. O. Ljungdahl, G. Mason-Buck, K. McGrath, C. Meydan, E. F. Mongodin, M. O. Moraes, N. Nagarajan, M. Nieto-Caballero, H. Noushmehr, M. Oliveira, S. Ossowski, O. O. Osuolale, O. Özcan, D. Paez-Espino, N. Rascovan, H. Richard, G. Rätsch, L. M. Schriml, T. Semmler, O. U. Sezerman, L. Shi, T. Shi, R. Siam, L. H. Song, H. Suzuki, D. S. Court, S. W. Tighe, X. Tong, K. I. Udekwu, J. A. Ugalde, B. Valentine, D. I. Vassilev, E. M. Vayndorf, T. P. Velavan, J. Wu, M. M. Zambrano, J. Zhu, S. Zhu, C. E. Mason, International MetaSUB Consortium, A global metagenomic map of urban microbiomes and antimicrobial resistance. Cell. 184, 3376–3393.e17 (2021).

21. N. S. Basrur, M. E. De Obaldia, T. Morita, M. Herre, R. K. von Heynitz, Y. N. Tsitohay, L. B. Vosshall, mutant male mosquitoes gain attraction to human odor. Elife. 9 (2020), doi:10.7554/eLife.63982.

22. O. J. Bosch, M. Geier, J. Boeckh, Contribution of fatty acids to olfactory host finding of female Aedes aegypti. Chem. Senses. 25, 323–330 (2000).

23. D. Lucas-Barbosa, M. DeGennaro, A. Mathis, N. O. Verhulst, Skin bacterial volatiles: propelling the future of vector control. Trends Parasitol. 38, 15–22 (2022).

24. S. Conlan, L. A. Mijares, NISC Comparative Sequencing Program, J. Becker, R. W. Blakesley, G. G. Bouffard, S. Brooks, H. Coleman, J. Gupta, N. Gurson, M. Park, B. Schmidt, P. J. Thomas, M. Otto, H. H. Kong, P. R. Murray, J. A. Segre, Staphylococcus epidermidis pan-genome sequence analysis reveals diversity of skin commensal and hospital infection-associated isolates. Genome Biol. 13, R64 (2012).

25. Y. E. Chen, D. Bousbaine, A. Veinbachs, K. Atabakhsh, A. Dimas, V. K. Yu, A. Zhao, N. J. Enright, K. Nagashima, Y. Belkaid, M. A. Fischbach, Engineered skin bacteria induce antitumor T cell responses against melanoma. Science. 380, 203–210 (2023).

